# Overexpression of AMPKγ2 increases AMPK signaling to augment human T cell metabolism and function

**DOI:** 10.1101/2022.10.01.510473

**Authors:** Erica L. Braverman, Margaret A. McQuaid, Darlene A. Monlish, Andrea K. Dobbs, Manda J Ramsey, Archana Ramgopal, Harrison Brown, Craig A. Byersdorfer

**Author notes:** Corresponding author: Dr. Craig A. Byersdorfer, Division of Blood and Marrow Transplant and Cellular Therapies, Rangos Research Building, 4401 Penn Ave, Pittsburgh, PA 15224.

## Abstract

T cell-based cellular therapies benefit from a product with reduced differentiation and enhanced oxidative metabolism. Methods to achieve this balance without negatively impacting T cell expansion or impairing T cell function have proven elusive. AMP-activated protein kinase (AMPK) is a cellular energy sensor which promotes mitochondrial health and improves oxidative metabolism. We hypothesized that increasing AMPK activity in human T cells would augment their oxidative capacity, creating an ideal product for adoptive cellular therapies. Lentiviral transduction of the regulatory AMPKγ2 subunit stably enhanced intrinsic AMPK signaling and promoted mitochondrial respiration with increased basal oxygen consumption rates (OCR), higher maximal OCR, and augmented spare respiratory capacity. These changes were accompanied by increased mitochondrial density and elevated expression of proteins involved in mitochondrial fusion. AMPKγ2-transduction also increased T cell glycolytic activity. This combination of metabolic reprogramming enhanced *in vitro* T cell expansion while promoting memory T cell yield. Finally, when activated under decreasing glucose conditions, AMPKγ2-transduced T cells maintained higher levels of both proliferation and inflammatory cytokine production. Together, these data suggest that augmenting intrinsic AMPK signaling via overexpression of AMPKγ2 can improve the expansion and function of human T cells for subsequent use in adoptive cellular therapies.

**Key points:** Lentiviral Transduction of AMPKγ2 increases oxidative metabolism in human T cells

AMPKγ2 transduction enhances *in vitro* proliferation without inducing exhaustion

AMPKγ2-transduced T cells function better under low glucose conditions

## INTRODUCTION

Significant recent attention has focused on T cell metabolism and its influence on T cell fate and function (1–3). Adoptive cellular therapies for cancer, in particular, have highlighted a role for immune cell metabolism, with chimeric antigen receptor (CAR) T cells and tumor-infiltrating lymphocytes (TILs) demonstrating reduced efficacy due to metabolic exhaustion and/or decreased persistence of the transferred cells (4–6). In contrast, adoptive T cell therapies which increase oxidative metabolism and preserve mitochondrial health demonstrate enhanced performance both *in vitro* and during subsequent transfer *in vivo* (7, 8). For this reason, extensive efforts have been made to improve the metabolic capacity of T cells. Several interventions to reduce T cell differentiation and minimize metabolic exhaustion have been reported, including nutrient restriction, inhibition of protein synthesis, expansion in the presence of exogenous cytokines (e.g. IL-7 or IL-15), blockade of mitochondrial fission, promotion of mitochondrial fusion, and upregulation of mitochondrial biogenesis (9–15). While multiple approaches have successfully enhanced oxidative metabolism, a major drawback to many of these methods is a sharp reduction in T cell proliferation, limiting clinical applicability. In addition, numerous pharmaceutical interventions are operational only during *in vitro* expansion, which may not provide long-lasting functional benefit once cells are transferred *in vivo*. For this reason, the search continues to identify and develop durable and effective approaches to optimizing T cell metabolic health while still preserving proliferation and limiting *in vitro* differentiation. Modulating immune cell metabolism through manipulation of the cellular energy sensor, AMP-activated protein kinase (AMPK), offers significant potential to achieve this goal.

AMPK detects decreasing intracellular energy stores through a rise in AMP and a concomitant decrease in ATP. When the overall AMP/ATP ratio increases, AMPK becomes activated and then phosphorylates downstream targets to block anabolic growth and promote catabolic metabolism and mitochondrial efficiency. AMPK is well known for facilitating mitochondrial health through mitophagy and mitochondrial fusion (16) as well as antagonizing mammalian target of rapamycin (mTOR) signaling (17). The end result of these actions is increased scavenging of reactive oxygen species (ROS), enhanced fatty acid oxidation, and blockade of fatty acid synthesis (16,18–24). While each of these pathways can independently facilitate *in vivo* health and potentiate anti-tumor efficacy, modulating AMPK signaling has the benefit of addressing all three pathways simultaneously. Furthermore, global knock-out of AMPK in three different mouse tumor models significantly decreased tumor cell killing (25), suggesting that AMPK is responsible for controlling pathways critical for maximal T cell function *in vivo*.

AMPK is a heterotrimeric protein complex consisting of α, β, and γ subunits, each of which has multiple isoforms with tissue-specific distribution(26). The α subunit houses the kinase activity and is active when phosphorylated on amino acid Thr172. Upstream activators of AMPK include liver kinase B-1 (LKB) and calcium influx, which signals through Calcium/calmodulin-dependent protein kinase kinase (CAMKK) to activate AMPK following T cell receptor (TCR) stimulation (27). Once phosphorylated, the α subunit phosphorylates downstream targets, including Unc51-like kinase 1 (ULK1), acetyl-coA carboxylase (ACC), and Peroxisome proliferator-activated receptor-gamma coactivator-1 α (PGC1α) (24, 28–30). In counterbalance, phosphatases, including the heterotrimer protein phosphatase 2A (PP2A), dephosphorylate AMPKα to downregulate its activity (31). As a further level of control, AMPKα dephosphorylation can be prevented by actions of the regulatory γ subunit, which protects the phosphorylated α domain and thereby preserves its kinase activity while also providing additional allosteric activation to the AMPK complex when bound to AMP (32).

Many pharmacologic agents increase AMPK activity, metformin being the most well-known. However, many of these interventions, including metformin, increase intracellular AMP/ATP ratios by blocking mitochondrial respiration, in effect starving the cell. The downside of this approach is creation of mitochondrial dysfunction, which is counterproductive to optimal T cell function. Other agonists, including 5-Aminoimidazole-4-carboxamide ribonucleotide (AICAR), function as AMP mimetics. In addition to being non-specific (reviewed in (33)), increased AMPK signaling following mimetic treatment again relies on the cell feeling inhibited, a process which again downregulates critical cellular pathways, including cell growth and protein synthesis. Furthermore, different downstream signaling networks can be engaged depending upon the setting in which AMPK is activated. These differences are thought to be related both to the upstream activator as well as to the isoform makeup of the activated AMPK heterotrimer (34). Together, these previous studies suggest that forcing cells to increase AMPK signaling through perceived or actual nutrient starvation may hinder immune responses. In contrast, a method to selectively increase AMPK activity, downstream of natural signals, and with the possibility *of in vivo* durability, would be more ideal.

T cells express two isoforms of the regulatory AMPKγ subunit-AMPKγ1 and AMPKγ2, with AMPKγ1 being more highly expressed (35). The AMPKγ2 isoform, however, has a higher affinity for ADP than AMPKγ1, suggesting that increased expression of AMPKγ2 might improve AMPK activity via heightened sensitivity of the heterotrimer to energetic flux (36). Furthermore, AMPKγ2 has a longer N-terminus, a feature hypothesized to increase protection of the phosphorylated AMPKα domain (26). Thus, in searching for a means to increase AMPK signaling, we elected to overexpress the regulatory subunit AMPKγ2 in primary human T cells. Here, we demonstrate that overexpression of AMPKγ2 increases AMPK activity, enhances mitochondrial mass, heightens spare respiratory capacity, and improves overall metabolic efficiency, endowing T cells with characteristics ideal for subsequent use in adoptive cellular therapies.

## RESULTS

### AMPKγ2 overexpression increases AMPK activity in primary human T cells

Previous reports have demonstrated that the AMPKγ2 domain has higher affinity for ADP than AMPKγ1 and has a longer n-terminus, allowing for increased protection of the phosphorylated AMPKα domain (26, 36). We therefore hypothesized that overexpressing AMPKγ2 in human T cells would lead to increased and prolonged AMPK activity. To test this idea, primary human T cells were stimulated with CD3/CD28 Dynabeads followed by transduction with lentiviral constructs containing the AMPKγ2 sequence downstream from an elongation factor 1 alpha (EF1a) promoter and upstream from either GFP or RQR8 (38) separated by a T2A linker (Fig. 1A). AMPKγ2 exists in 5 defined isoforms (https://www.ncbi.nlm.nih.gov/gene/51422); for these experiments, we elected to overexpress isoform C, a naturally occurring sequence slightly shorter than the more actively produced isoform, isoform A, and one which allows for easy identification of transduced cells by western blot. Control cells were transduced with T2A-GFP or T2A-RQR8 “Empty” constructs. On day 5 post-transduction, GFP or RQR8 expression was assessed by flow cytometry and revealed efficient transduction of both plasmids (Fig. 1B). RNA harvested on Day 9 revealed a 5-fold increase in AMPKγ2 levels compared to Empty controls (Fig. 1C). As noted, the smaller isoform C could easily be separated from the naturally more abundant AMPKγ2 isoform A by immunoblot analysis, which confirmed increased isoform C protein expression in AMPKγ2-transduced cells (Fig. 1D). We then asked whether overexpression of the regulatory γ2 subunit was sufficient to increase AMPK activity. GFP+ CD4+ and CD8+ T cells were flow sorted on Day 9 and Thr172 phosphorylation of AMPKα assessed by immunoblot. Overexpression of AMPKγ2 increased phosphorylation of AMPKα in both CD4 and CD8 T cells, with AMPK activity further confirmed through phosphorylation of two well-known AMPK targets, Ser79 on Acetyl-CoA Carboxylase (ACC) and Ser555 on Unc-51 like autophagy activating kinase (ULK1) (Fig. 1E).

**Figure 1.**
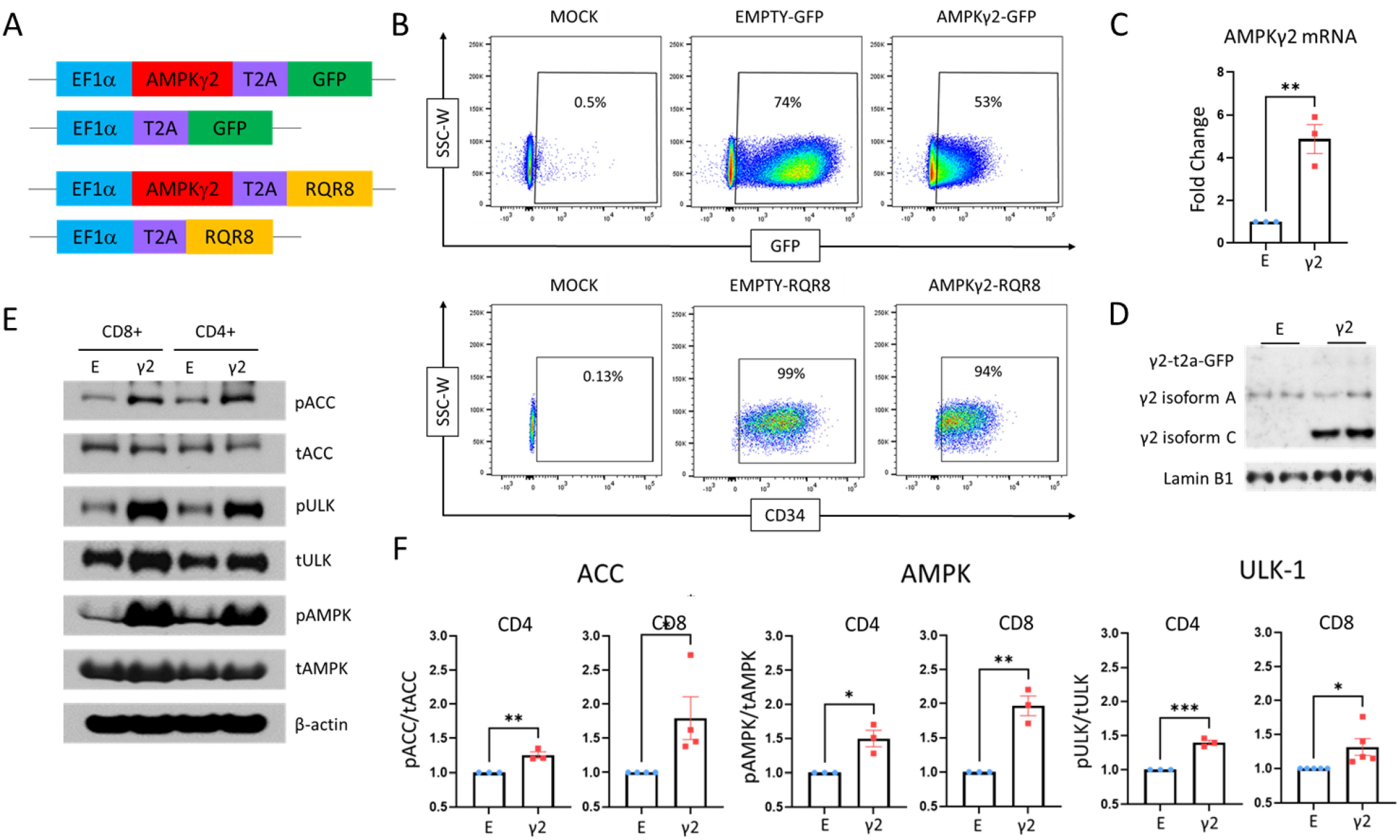
AMPKγ2 overexpression increases AMPK activity in human T cells. **(A)** Schematic of AMPKγ2 and GFP-only “Empty” control vectors with an EF1α promoter and either GFP or RQR8 expression tag. **(B-D)** Primary human T cells were mock transduced or transduced with AMPKγ2 or Empty plasmids. Expression was verified by flow cytometry for GFP or the CD34 motif of the RQR8 construct (B), fold-change in AMPKγ2 mRNA expression using qRT-PCR **(C),** and immunoblot to detect protein expression of AMPKγ2 (D). To note, studies utilized a shorter isoform of AMPKγ2 (isoform C), with lower expression levels at baseline, allowing for easy identification of a distinct, lower molecular weight band. (E) Human T cells were transduced with AMPKγ2-versus Empty constructs, cells lysates collected on days 9-12, and phosphorylation of AMPKα Thr172 (to detect AMPK activation), ACC Ser79, and ULK-1 Ser555 measured by immunoblot. (F) Densitometry was measured on immunoblots from multiple donors using ImageJ software. Values for AMPKγ2-transduced cells in each sample were normalized to Empty controls. All data represent 3 or more independent human donor samples. *p<0.05, **p<0.01, ***p<0.001 by paired Student’s T test.

### AMPKγ2 overexpression enhances metabolic capacity

We next sought to study the metabolic effects of AMPKγ2 overexpression. AMPKγ2- or Empty-transduced cells were expanded *in vitro* in the presence of IL-2 through day 9, then placed into a Seahorse Metabolic Analyzer to measure basal oxygen consumption rates (OCR), maximal respiration, and spare respiratory capacity (SRC). AMPKγ2-transduced cells increased all three parameters compared to Empty-transduced controls (Fig. 2A). We next determined whether these changes continued following TCR activation. A subset of day 9 cells were stimulated overnight with anti-CD3/CD28 Dynabeads followed by subsequent analysis of metabolic activity. Following stimulation, increases in basal OCR, maximal OCR, and SRC were even more pronounced. To investigate the etiology of this increased respiratory capacity, mitochondrial density was assessed using MitoTracker Red, which showed a reproducible and statistically significant increase in mitochondrial mass in AMPKγ2-transduced cells (Fig. 2B). These results were supported in CD8 T cells by a notable increase in PGC1α expression, a transcriptional co-activator known to promote mitochondrial biogenesis downstream of AMPK (28) (Fig. 2C).

**Figure 2.**
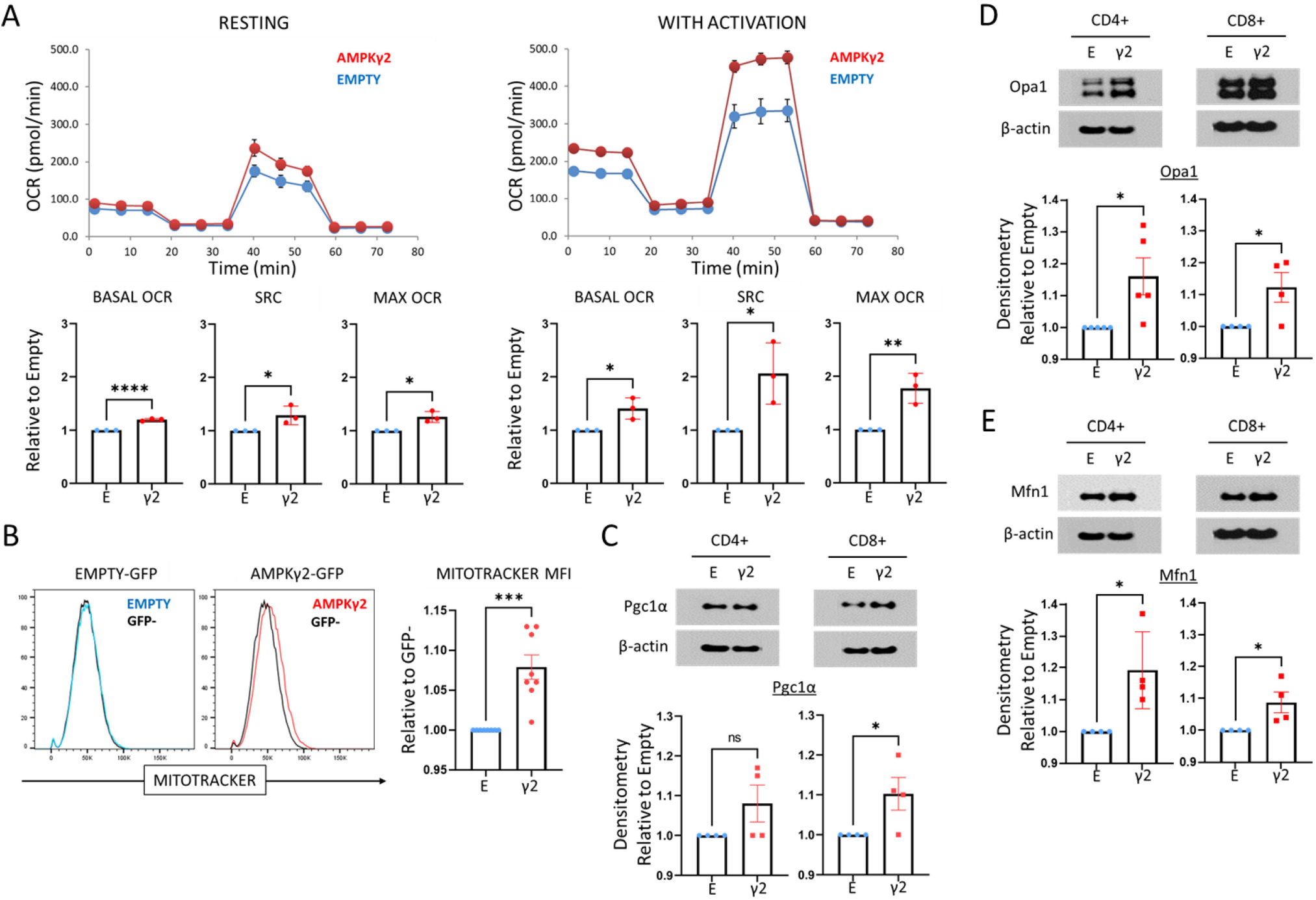
AMPKγ2 transduced Tcells enhance oxidative metabolism. **(A)** Human T cells were transduced with AMPKγ2 or Empty lentiviral vectors and expanded in IL-2. On day 9, a portion of cells were stimulated overnight with anti-CD3/CD28 Dynabeads and the next day both resting (left) and activated (right) cells were assessed for oxidative capacity utilizing the Seahorse Metabolic Analyzer. Bar graphs represent data from 3 individual human donors. (B) AMPKγ2- and Empty-transduced T cells were assessed for mitochondrial density utilizing MitoTracker Red (left). Fluorescence was compared to GFP^neg^ controls within each sample to standardize staining between groups. Differences in median fluorescence intensity (MFI) between GFP+ and GFP^neg^ cells was normalized to differences seen in the Empty control from 5 human donors (right). **(C-E)** GFP+ AMPK-versus Empty-transduced T cells were sorted into CD4+ and CD8+ subsets on Day 9 post-stimulation and protein expression of the transcriptional co-activator PGClα **(C)** or mitochondrial fusion proteins OPA1 **(D)** and MFN1 **(E)** assessed by immunoblot. Bar graphs represent data from 3-4 human donors. *p<0.05, **p<0.01, and ***p<0.001, ****p<0.0001 by paired Student’s T test.

Given that both the morphology and density of mitochondria affect their efficiency, we quantitated expression of OPA1 and MFN1, two mitochondrial fusion proteins whose expression has been shown to increase in response to enhanced AMPK activity (16). Indeed, AMPKγ2 transduction upregulated expression of OPA1 and MFN1 in both CD4+ and CD8+ cells (Fig. 2D,E), suggesting that both a greater mitochondrial density and increased mitochondrial fusion may be contributing to the observed increase in oxidative capacity.

### AMPKγ2 overexpression enhances memory T cell yield

We next sought to characterize whether AMPKγ2 transduction impacted T cell differentiation. This feature was of particular interest given that adoptively transferred T cells which retain a more memory-like phenotype function more effectively *in vivo* (13,14, 39, 40). AMPKγ2 transduction increased the percentage of central memory-like T cells on Day 9 of culture as measured by increased co-expression of CD62L/CCR7 (Fig. 3A-B). Further, to assess whether this AMPK effect was additive or irrelevant in the face of “memory inducing” cytokines, we transduced T cells in IL2, followed by subsequent culture in IL7 and IL15, a process shown to promote central memory cells (14, 41, 42). Again, AMPKγ2 transduction increased the yield of central memory-like T cells on Day 9 of culture (Fig. 3C-D). Interestingly, while the increase in memory CD8+ T cells was similar in IL2 versus IL7/IL15, the increase in CD4+ memory T cells was significantly more dramatic with IL7/IL15 (Fig. 3E), suggesting a potential synergistic effect between these cytokines and AMPK on the development of CD4+ memory T cells.

**Figure 3.**
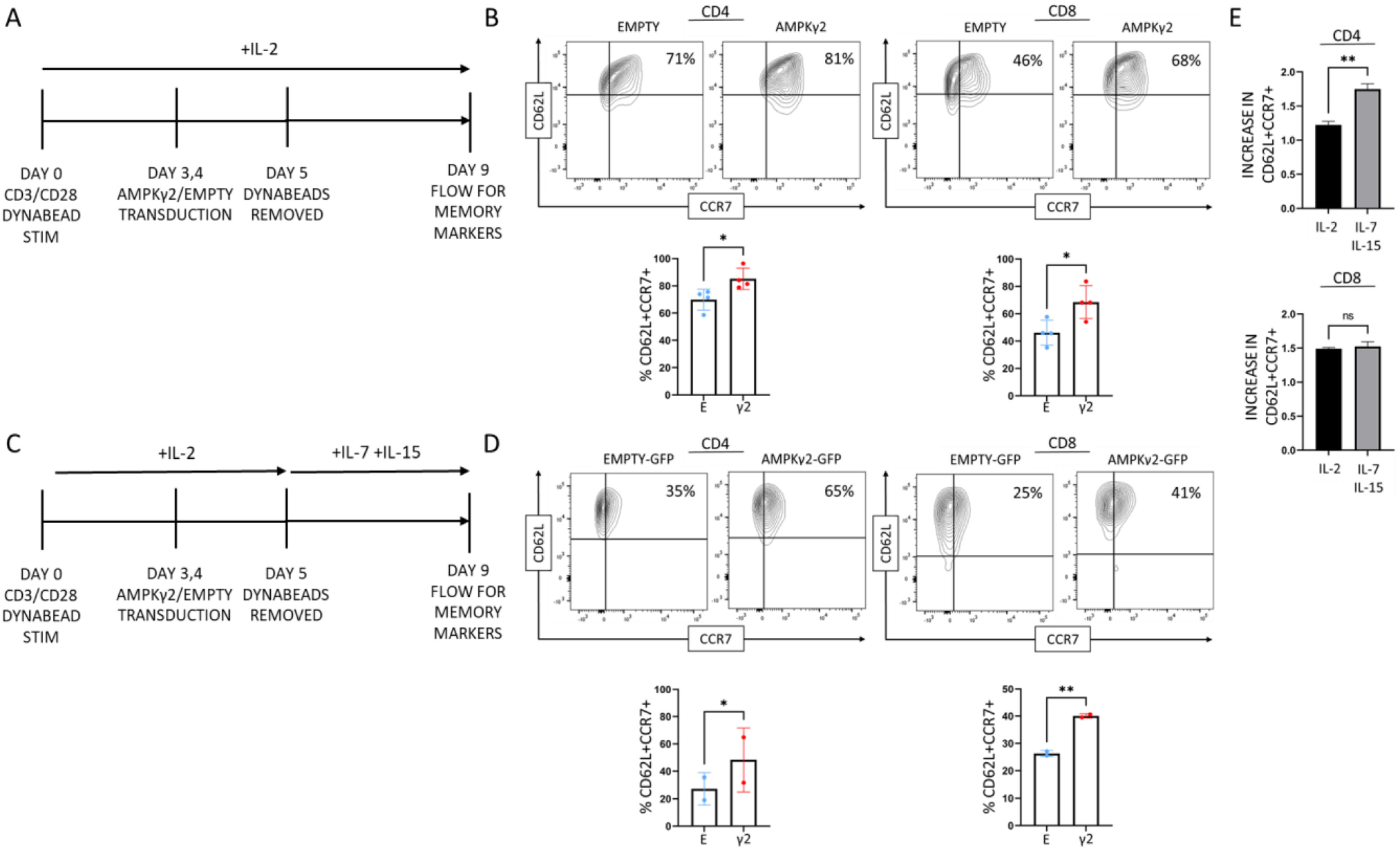
AMPKγ2 transduction enhances yield of CD4 and CD8 memory T cells. **(A-B)** AMPKγ2- and Empty-transduced cells were expanded *in vitro* in the presence of IL-2 and assessed on Day 9 for cell surface expression of CD62L and CCR7 to identify central memory-like T cells. **(C-D)** A second group of cells were stimulated in IL-2 and then expanded in IL-7 and IL-15 until Day 9, when they were stained for cell surface expression of CD62L and CCR7. (E) Relative increases in CD62L+CCR7+ cells were calculated based upon the ratio of double positive AMPK-transduced to Empty-transduced controls. Bar graphs for B, D represent data from 2-4 individual human donors. *p<0.05 and **p<0.01 by paired Student’s T test.

**Figure 4.**
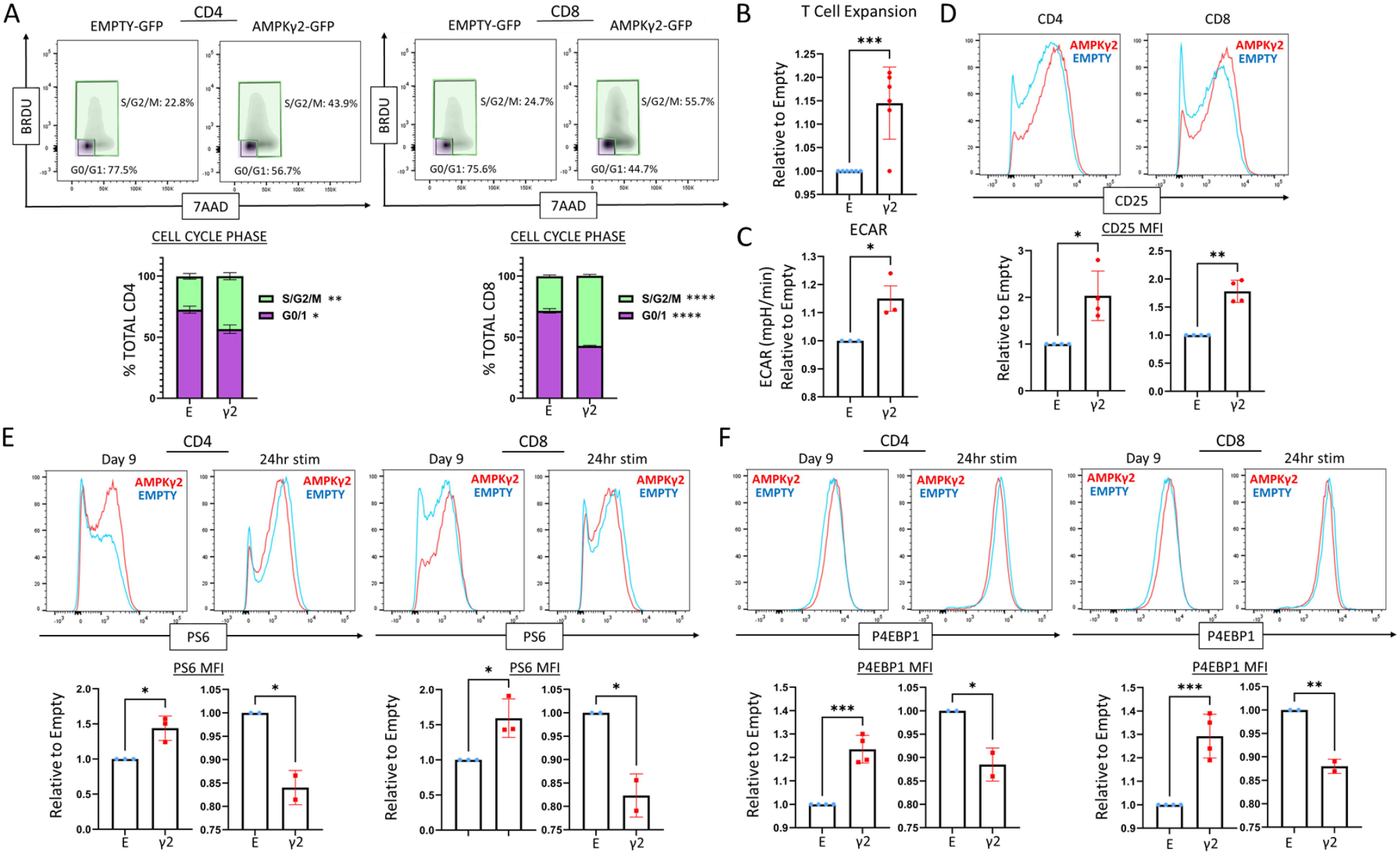
AMPK activation increases T cell expansion, cell cycling, and mTOR activity. **(A)** AMPK- and Empty-transduced T cells were expanded *in vitro* for 7-9 days, incubated with BrdU for the final 2 hours, and co-stained with 7AAD. Plots are divided into resting (G0/G1) and cycling (S, G2/M) phases. Graphs represent data from 3 human donors. (B) AMPKγ2-transduced human T cells (GFP+) were manually counted between Day 5 and 7 to calculate the doubling rate per 24 hours, which was then normalized to the Empty-transduced control for each donor. Graph represents data from 6 independent human donors. **(C)** Transduced cells were harvested on Day 9 of culture and stained for CD25. MFI values were compared between multiple donors. (D) Transduced human T cells were assessed on day 9 for baseline extracellular acidification rates (ECAR) using the Seahorse Metabolic Analyzer. **(E-F)** Empty- and AMPKγ2-transduced T cells were expanded until day 9, followed by assessment of mTOR activity using antibodies against phosphorylated S6 **(E)** and 4EBP1 **(F).** Cells were assessed either at rest (day 9) or following 24 hours of CD3/CD28 stimulation (24hr stim). The MFI for *P-S6 or *P-4EBP1 from each donor was normalized to expression in Empty-transduced controls and this value was then compared between donors. Bar graphs represent data from 4 human donors.*p<0.05, **p<0.01, ***p<0.001, and ****p<0.0001 by paired Student’s T test.

### AMPK-transduction boosts *in vitro* T cell proliferation

Many methods to increase oxidative metabolism (e.g. inhibiting glycolysis) have also successfully increased central memory T cell pools but end up restricting the expansion of treated cells (12). This drawback has limited the clinical utility of such approaches due to the need for sufficient *in vitro* expansion to allow for clinical administration. We therefore sought to examine how increased AMPK activity impacted cellular expansion *in vitro*. To assess cell cycling, we measured BrdU uptake between days 7 and 9 of culture followed by counterstaining with 7-AAD. Consistently, both CD4+ and CD8+ AMPKγ2-transduced T cells demonstrated fewer resting G0/G1 cells and more cells in the S/G2/M phases of the cell cycle (Fig. 3A). Further, quantitation of GFP+ cell numbers on day 5 and again on day 7 demonstrated a statistically significant increase in the doubling rate of AMPKγ2-transduced cells over a 24-hour period (Fig. 3B). This increase in cell cycling was accompanied by increased expression of CD25, the α chain of the IL2 receptor, suggesting that AMPKγ2-transduced cells may be more sensitive to levels of IL-2 in the immediate environment (Fig. 3C). AMPK can also promote glycolysis by increasing glucose uptake and mediating downstream effects on phosphofructokinase-1 and phosphorylation of 6-phosphofructo-2-kinase (43, 44). In support of this idea, the extracellular acidification rate (ECAR), an indirect measure of glycolysis, also significantly increased AMPKγ2-transduced T cells (Fig. 4D).

Highly proliferative cells also rely on increased signaling through mTOR to provide the necessary building blocks for continued division. Given the known antagonism between the AMPK and mTOR pathways, we questioned whether mTOR activity was affected by AMPKγ2 transduction. We found that mTOR signaling, as assessed by phosphorylation of downstream targets S6 and 4EBP1, increased in AMPKγ2-transduced cells on Day 9 of culture, correlating with their ongoing increased proliferation. However, 24 hours after re-stimulation, mTOR signaling slightly decreased in AMPK-transduced cells, consistent with an acute role of AMPK in decreasing mTOR signaling (45) (Fig. 4E-F). Therefore, AMPK-transduced cells maintain their proliferation *in vitro*, which correlates with increased glycolysis as well as a relative increase in mTOR signaling at later timepoints.

### AMPK-transduction does not accelerate exhaustion

Given the increased proliferation and enhanced glycolytic rate of AMPKγ2-transduced cells, we worried about the possibility of T cell exhaustion. To address this point, we examined transduced cells for the expression of activation/exhaustion markers PD1, Tim3, and LAG3. Reassuringly, there was no significant difference in PD1, Tim3, or LAG3 expression in AMPKγ2-transduced versus control T cells on day 9, nor did AMPK-transduced cells demonstrate an inability to upregulate these activation markers upon re-stimulation, supporting the idea that AMPKγ2-transduced cells were neither exhausted nor functionally impaired (Fig 5A). While the expression of phenotypic markers was reassuring, we also placed AMPKγ2-versus Empty-transduced into an *in vitro* exhaustion protocol (46, 47) (Fig. 5B). Expanded cells were re-stimulated with CD3/CD28 Dynabeads every 48 hours over 6 days, with cell counts, IL2, and IFNG measured at each timepoint. Cells were then placed into a mixed leukocyte reaction (MLR) with irradiated, allogeneic non T cells and examined for proliferation by BRDU uptake after 72 hours, with supernatant cytokine production measured by LEGENDPLEX analysis. During CD3/CD28 re-stimulation, AMPKγ2-transduced cells showed no deficiency in cell count or cytokine production and in fact made more IFNγ upon initial re-stimulation (Fig. 5C). In the MLR, AMPK-transduced cells had enhanced proliferative capacity as measured by BRDU uptake, with increased production of inflammatory cytokines IL-8 and IL-6 (Fig. 5D-E). Thus, despite increased proliferation and glycolysis, AMPK-activated cells remained more memory-like, did not overexpress exhaustion markers, and were more inflammatory upon reactivation, even after repeat stimulation *in vitro*.

**Figure 5.**
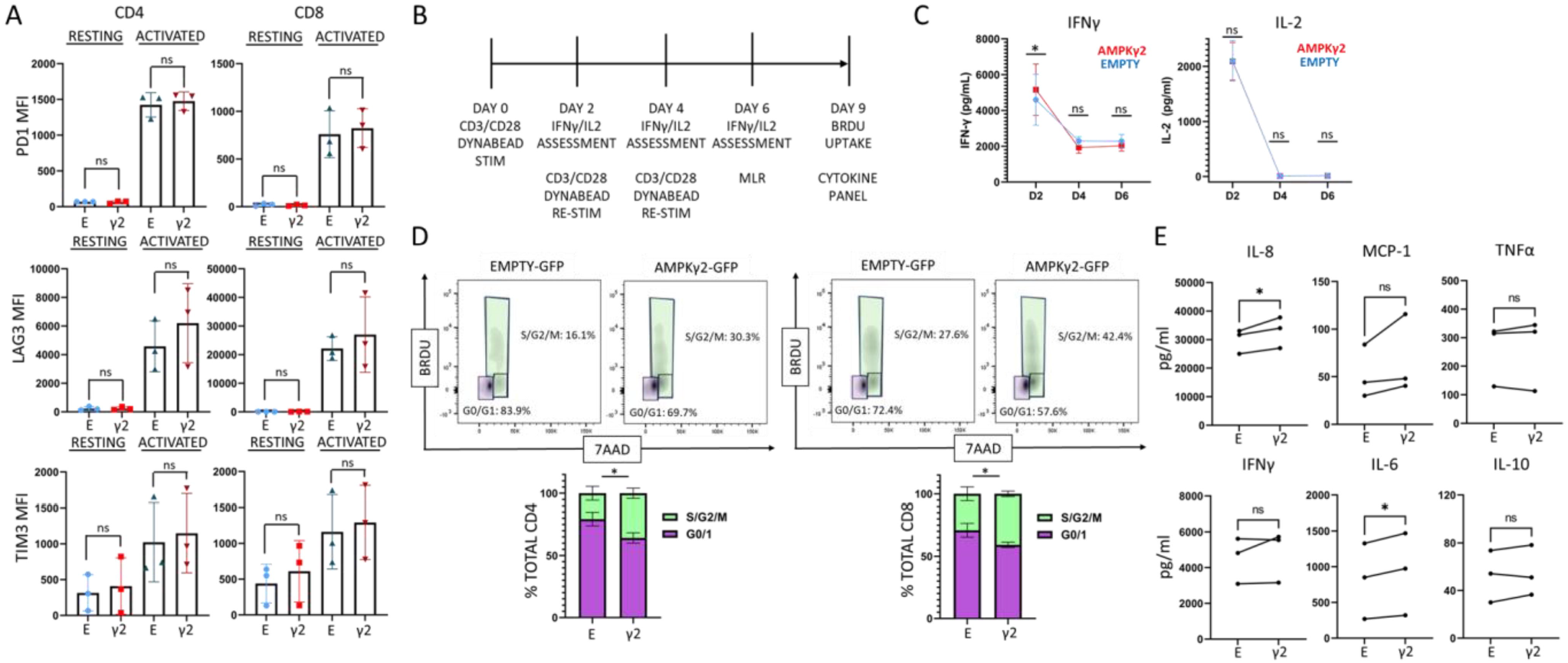
AMPK-transduction does not accelerate exhaustion. **(A)** AMPKγ2- and Empty-transduced cells were expanded *in vitro* in the presence of IL-2 and assessed on Day 9 for cell surface expression of PD-1, Tim-3, and Lag-3 (resting). A portion of cells were then re-stimulated with CD3/CD28 Dynabeads for 24 hours and reassessed for PD-1, Tim-3, and Lag-3 expression (Activated). (B) A timeline of the *in vitro* exhaustion protocol, with **(C)** plots representing IFNγ and IL-2 production as measured by ELISA. **(D-E)** After three consecutive re-stimulations, T cells were placed with allogeneic APCs for 72 hours, incubated with BrDU for two additional hours, counterstained with 7AAD, and measured for cell cycle analysis by flow cytometry (D). Media was also harvested from MLR cultures at 72 hours and assessed for differences in cytokine expression between AMPK- and Empty-transduced T cells using LEGENDplex analysis (E). Data represent 3 human donors. *p<0.05 by paired Student’s T test.

### Increased AMPK activity improves T cell function during glucose restriction

While the enhanced performance of AMPK-transduced cells following repeated *in vitro* stimulation was encouraging, we wanted to further understand if AMPK-driven improvements in metabolic capacity translated to improved function under nutrient-limiting stress. To this end, we activated AMPKγ2- or Empty-transduced cells with T cell TransAct (Miltenyi) in either standard RPMI, containing 11mM glucose (high glucose (HG)), or in RPMI with glucose reduced to 5.5 mM (physiologic glucose (PG)). Proliferative capacity was assessed at 72 hours. While AMPKγ2-transduced cells showed no differences in cell cycling under HG conditions (Supp. Fig. 1A), AMPKγ2 cells stimulated in PG had fewer cells in G0/G1 and more cells in the S/G2/M phases of the cell cycle compared to control cells (Fig. 6A-B). To assess whether this advantage persisted following a more physiologic activation signal, day 9 cells were stimulated with allogeneic antigen presenting cells (APCs) in a one-way MLR and cytokine production measured from MLR supernatants utilizing LEGENDplex analysis. While there were no significant differences in HG (Supp. Fig. 1B), levels of MCP-1,1L-8, and TNF increased following allogeneic activation of AMPKγ2- transduced cells in PG. Other pro-inflammatory cytokines including IFNγ and IL-10 trended towards increased expression (Fig. 6C). Further, in comparing cytokine production between HG and PG conditions, AMPKγ2-transduced cells actually increased production of MCP-1, TNFα, IL-6, and IL-10 in PG, whereas Empty-transduced controls exhibited no change in cytokine levels based upon glucose content (Supp. Fig. 1C). At the same time, Empty-transduced controls produced less IL-8 and IFNγ in PG while AMPKγ2-transduced cells maintained levels of these cytokines (Supp. Fig. 1D). Given the improved function of AMPKγ2-transduced cells in PG, we chose to further assess their function under true glucose tension (2.25mM), which we refer to as “low glucose” (LG) (Fig. 6D). In this setting, AMPK-transduced cells continued to demonstrate both enhanced cell cycling (Fig. 6E) and increased inflammatory cytokine production (Fig. 6F). These data imply that AMPKγ2-transduced cells maintain inflammatory cytokine production in spite of glucose restriction and together support an ability of AMPKγ2-transduced cells to increase both proliferation and effector functions when glucose is limited.

**Figure 6.**
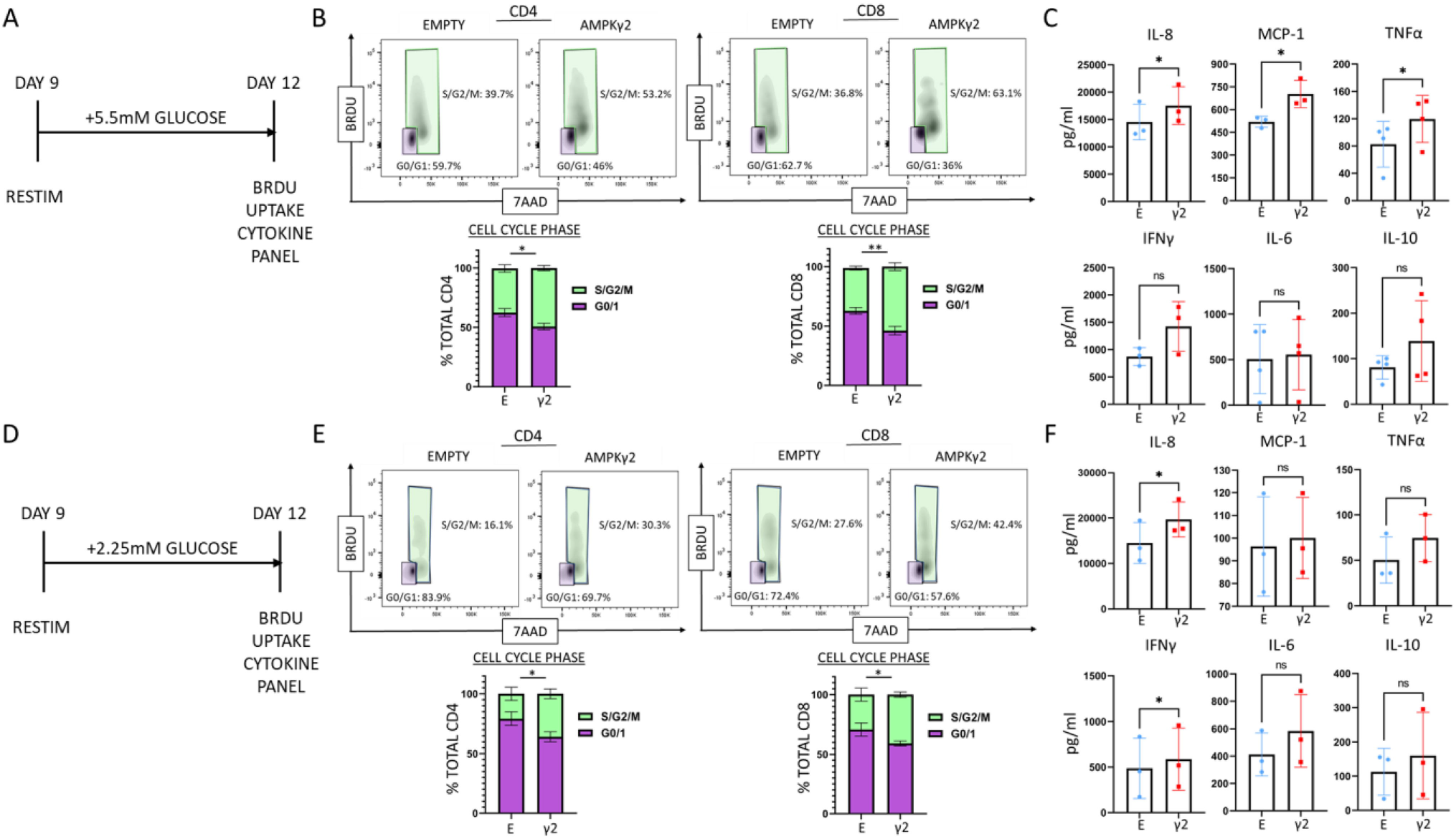
AMPKγ2 transduced cells exhibit improved function during nutrient restriction. AMPK- and Empty-transduced T cells were cultured for up to 12 days, re-stimulated in 5.5 mM (physiologic) (A-B) or 2.25 mM (low) (D-E) glucose for 72 hours, incubated with BrdU for 2 additional hours, counter-stained with 7AAD, and assessed for cell cycling. **(C, F)** AMPK- and Empty-transduced T cells were cultured for 9-12 days before being re-plated 1:1 with allogeneic APCs in either **(C)** physiologic (5.5 mM) or **(F)** low (2.25mM) glucose for an additional 72 hours. Media was harvested and differences in cytokine expression assessed between AMPK- and Empty-transduced T cells using LEGENDplex. Bar graphs represent data from 3-4 independent donors. *p<0.05 and **p<0.01 by paired Student’s T test.

## DISCUSSION

Adoptive cellular therapies, including CAR-T cells, are often limited by terminal differentiation of the cellular product and metabolic exhaustion of the transferred cells. These less desirable phenotypes result from a number of secondary causes including glycolysis-inducing *in vitro* expansion protocols, the immunosuppressive and nutrient-deplete *in vivo* tumor environment, and the exhausted status of cells when initially harvested from patients. Methods to ameliorate this metabolic and functional insufficiency are sorely needed to improve and expand the use of adoptive cellular therapies. One unique advantage of *in vitro* expansion is the opportunity to alter T cell properties and improve their subsequent function prior to return to patients. Manipulating a variety of metabolic pathways has been attempted, but alterations have thus far seen limited clinical application due, in part, to restricted expansion resulting from the methods used to promote oxidative metabolism (12).

In our model, lentiviral transduction of the regulatory AMPKγ2 subunit increased activity of the AMPK heterotrimer, a major regulator of oxidative metabolism and guardian of mitochondrial health. Notably, our data suggest that increasing AMPK signaling via AMPKγ2 overexpression enhances oxidative metabolism while still increasing the *in vitro* expansion of transduced cells. Importantly, the enhanced respiration seen in resting AMPKγ2-transduced cells increased further following subsequent TCR activation. This powerful finding suggests that metabolically reprogramming the AMPK pathway is likely to provide metabolic benefits during T cell stimulation, where underperforming cells require increased metabolic activity the most. We further found that activation of AMPK increased the yield of central memory T cells, even in the setting of “memory-inducing” cytokines IL7 and IL15, consistent with the fact that AMPK is known to be more active in memory than effector T cells (48). More interesting, perhaps, was the apparent synergy between IL7/IL15 and AMPK in augmenting CD4+ central memory T cell generation. This finding deserves further inspection, particularly given recent data suggesting that long-term cures in leukemia patients treated with CART therapy correlate with the size of the pool of long-term CD4+ memory T cells (49).

AMPKγ2-transduced T cells also outperformed control cells in both proliferation and pro-inflammatory cytokine production when activated under physiologic and low glucose conditions, suggesting that transduced cells would have an advantage in glucose-restricted settings *in vivo* (50). The fact that these changes were not seen in the setting of high glucose is supported by other studies in which the advantages of increased AMPK signaling required cellular stress to translate into functional differences (51). In evaluating the mechanism underlying these improvements, we found increases in proteins related to both mitochondrial biogenesis and mitochondrial fusion, consistent with known effects of AMPK signaling (1,16). These mitochondrially-centered changes also likely contribute to the notable increase in SRC, which has been a hallmark of T cells capable of improved functional ability *in vivo* (1, 7). In addition, upregulation of ULK1 signaling and its effect on mitophagy may further contribute to overall mitochondrial health and improved oxidative efficiency (16, 52).

In addition to the expected changes in oxidative metabolism, AMPKγ2-overexpressing T cells also enhanced extracellular acidification, suggesting increased glycolysis. This effect is consistent with previous reports where AMPK signaling increased glycolysis alongside increased expression of GLUT transporters (53) and enhanced phosphorylation of 6-phosphofructo-2-kinase (20, 43). Increased flux through the proximal steps of glycolysis could also create more pyruvate to shuttle to the TCA, feeding an increase in oxidative metabolism (50). Elevated glycolysis is also in line with the increased growth and proliferation seen in AMPKγ2-transduced cells, where more cells are found within active phases of the cell cycle. Our data on CD25 expression suggests that at least a portion of this increased proliferation may be secondary to a heightened responsiveness to IL2, a point we intend to explore further. Notably, despite increased *in vitro* expansion, there was no evidence of impending exhaustion in AMPKγ2- transduced cells as measured by PD1, TIM3, and LAG3 expression and no impairment in upregulation of these markers upon further TCR stimulation. Further, AMPKγ2-transduced cells outperformed controls following a program *of in vitro* exhaustion - supporting the view that AMPKγ2-transduced cells are capable of full activation despite prolonged activity *in vitro*.

Increased cell proliferation also correlated with increased mTOR activity, a finding contradictory to the traditionally accepted antagonism between AMPK and mTOR signaling (17, 54). However, when mTOR activity was evaluated immediately following TCR stimulation, it was universally lower in AMPKγ2- transduced cells, demonstrating the expected counterbalance between these two pathways. One hypothesis is that persistent AMPK signaling creates a more metabolically efficient cell through promotion of mitochondrial health. Because of this ongoing efficiency, AMPKγ2-transduced cells maintain a higher level of proliferation as they expand in culture, with subsequent, compensatory increases in mTOR signaling. Another possible explanation concerns the function of AMPKγ2 relative to AMPKγ1. It is known that AMPKγ1 and AMPKγ2 isoforms function differently with respect to when they are activated, their intracellular localization, and the subsequent pathways they activate (26, 55). Importantly, AMPKγ2 is activated earlier in energy depletion than AMPKγ1 due to its higher affinity for ADP, potentially influencing pathways meant to prepare cells for pending energetic stress, including those charged with improving metabolic efficiency and energetic plasticity. In contrast, pathways which shut down anabolic growth, such as blocking mTOR signaling, are more heavily downstream of AMPKγ1, including the cellular response to metformin (56). Given that T cells more highly express AMPKγ1 at baseline, switching to prominent AMPKγ2 signaling may undermine traditional AMPK signaling pathways and instead promote the pro-inflammatory and proliferative phenotype seen in the AMPKγ2 overexpressing cells. AMPK is also activated downstream of the TCR, where it tailors T cell responses to energy availability (20). Because our lentiviral method augments endogenous AMPK signaling without relying on cellular starvation, our approach may allow for a deeper understanding of the connection between AMPK signaling and TCR-driven activation and growth. Indeed, such a connection may help explain why memory T cells, which have increased AMPK activity, show enhanced proliferation and more efficient cytokine generation upon re-activation (57).

A hypothetical concern when beginning these studies was that AMPKγ2-transduced cells would enforce catabolic pathways and simultaneously block cellular growth during nutrient restriction, a disadvantageous scenario for adoptively transferred cells. Reassuringly, cells transduced with AMPKγ2 outperformed control cells when glucose concentrations were restricted, implying that even in the setting of limited external energy sources, AMPKγ2-transduced cells can still overcome nutrient limitations to generate sufficient ATP to allow for increased function. This capability may reflect the role of AMPKγ2 in favoring metabolic plasticity over anabolic blockade, though may also be a consequence of enhanced mitochondrial biogenesis and fusion with more efficient energy production. Regardless of the cause, an ability to maintain enhanced function in the setting of limited glucose makes increasing AMPK activity through AMPKγ2 overexpression a particularly attractive method to improve cellular therapies *in vivo*.

Based on a wealth of data from the literature, T cells with increased oxidative metabolism and limited *in vitro* differentiation are expected to persist longer and perform better *in vivo* (10,12, 14, 15). Thus, the next step will be to trial AMPKγ2 transduction in a variety of cellular therapies and animal models to assess for the positive effects of enhanced AMPK activity on *in vivo* cellular functions. It could be envisioned that many cell-based therapies, including the metabolic reprogramming of TIL or the reshaping of an anti-viral immune response, may also benefit from an intervention that not only improves T cell growth and viability *in vitro*, but also potently enhances inflammatory functions in nutrient-deplete microenvironments. From the standpoint of feasibility, many therapies, including CAR T cell generation, already require lenti- or retroviral transduction coupled with the additional advantage that genetic manipulation is transferred with the T cells *in vivo*, potentially enhancing efficacy long after transfer.

In summary, increasing AMPK activity in primary human T cells through lentiviral-driven overexpression of AMPKγ2 upregulates oxidative metabolism, improves central memory formation, and improves *in vitro* expansion. Together, these characteristics provide an attractive method to improve current adoptive cellular therapies. Further, this work suggests a potentially exciting role for isoform specific control of AMPK signaling, the understanding of which could lead to an increased ability to further manipulate a master metabolic regulator towards therapeutic gain. Work is ongoing to both further uncover the role of specific AMPKγ isoforms in AMPK signaling in T cells, as well as to assess the potential contribution of AMPKγ2 overexpression to multiple T cell-based therapies *in vivo*.

## Acknowledgements

The authors would like to thank Linda McAllister-Lucas and Kelly Bailey for their careful review of this manuscript.

## MATERIALS AND METHODS

### Virus Production

The AMPKγ2 sequence was amplified from a commercially available plasmid (Addgene #23689) and cloned into either a pSICO (Addgene #31847) or pHR (similar to Addgene #14858, kind gift from Jason Lohmueller, UPMC Hillman Cancer Center) backbone, followed by addition of a T2A linker and either GFP or RQR8 tag (38). Transformed bacterial cultures were grown overnight in Terrific Broth (Sigma Aldrich) and plasmids isolated using QIAGEN QIAmp Miniprep Plasmid Isolation Kit 250. HEK293Ts (ATCC) were cultured in DMEM media (Gibco #11966-025) containing 10% FBS, Pen Strep, 2mM L-Glutamine, and MEM Non-Essential Amino Acids. Early passage cells were transfected using an Invitrogen Lipofectamine 3000 Transfection kit with 2500ng of RSV-REV, PMD-2G, and PRRE plasmid and 10,000ng of either AMPKγ2-t2a-GFP/RQR8 or t2a-GFP/RQR8 “empty” plasmid. After 24 hours, supernatant was replaced with IMDM media (Gibco #12440-053) containing 10% FBS. Supernatant containing viral particles was harvested at 48 and 72 hours, combined with Lenti-Pac (GeneCopoeia), and incubated at 4 degrees C overnight. Viral supernatants were then centrifuged at 3500x (g) for 25 minutes at 4 degrees, resuspended in DMEM, and either frozen at −80C or used immediately.

### T cell isolation, transduction, and culture

De-identified buffy coats were obtained from healthy human donors (Vitalant), diluted with PBS, layered over lymphocyte separation medium (MPbio), and centrifuged at 400 xg and 25 degrees for 20 minutes with no brake. The PBMC layer was removed and T cells isolated using the Miltenyi Biotec Human Pan T cell isolation kit. Purified T cells were resuspended in AIM-V +5% SR (Gibco #A25961-01) and plated with Dynabeads™ Human T-Activator CD3/CD28 for T Cell Expansion and Activation (ThermoFisher) at a 2:1 ratio for 48 hours. Transduction was then performed per manufacturer’s instructions utilizing retronectin coated plates (Takara). Cells were removed from Dynabeads by magnetic separation on Day 5 post-stimulation and expanded in AIM-V media with 5% SR containing IL-2 at 100IU/ml, or with IL-7 and IL15 at 20ng/ml. Further assessments were performed between Day 9 and 12 of culture. For restimulation experiments, cells were re-plated with Dynabeads at 1:1 ratio for up to 72 hours.

### Protein Isolation and Immunoblot

CD8+ and CD4+ T cells were first separated on day 9 utilizing the STEMCELL CD8 positive selection kit according to manufacturer’s instructions and plated overnight in AIM-V + 5% SR. The following day, GFP+ cells were flow sorted directly into 10% trichloroacetic acid (TCA) and lysates centrifuged at 16,000x(g) at 4*C for ten minutes, washed twice in ice cold acetone, resuspended in solubilization buffer (9M Urea containing 1% DTT and 2%Triton X and NuPAGE lithium dodecyl sulfate sample buffer 4X (Invitrogen) at a 3:1 ratio), and heated at 70*C for 10 minutes (58). Protein gel electrophoresis was performed on ice using NuPAGE 4-12% Bis-Tris Protein Gels (Invitrogen) at 135V. In some cases, protein samples were heated to 95C for 5 minutes prior to gel loading. Protein was transferred to Invitrolon™ 0.45μm PVDF membranes (Invitrogen) at 30V on ice for one hour. Membranes were blocked in Tris Buffered Saline-Triton containing 5% nonfat milk and immunoblotting performed according to the Cell Signaling Technologies Western Blot Protocol. Blots were stripped for 10 minutes (Restore PLUS Western Blot Stripping Buffer, Thermo scientific) prior to re-probing. Antibodies used for immunoblotting are listed in supplemental Table 1. Blots were developed with Super Signal West Femto chemiluminescence reagents (Thermo Fisher Scientific), detected by CL-X Posure Film (Thermo Scientific), and scanned in grayscale with an Epson V600 scanner. Images were cropped using ImageJ Software (version 1.47T), inverted, and densitometry quantitated in an area encompassing the largest band, followed by quantitation of all subsequent bands using the same 2-dimensional area.

### Flow Cytometry

Cells were washed with PBS + 2% fetal bovine serum (FBS) before staining with antibodies at 1:100 dilution for 30 minutes. For intracellular stains, cells were fixed per manufacturer’s instructions using Fix/Perm kit (Cat #:88-8824-00, Invitrogen) and then stained with antibodies at 1:100 dilution. Antibodies and other flow cytometry reagents are listed in Supplemental Table 2. MitoTracker Red (Invitrogen) staining was performed at 1 nM in PBS with incubation at 37 degrees for 15 minutes. CellRox (Invitrogen) staining was performed in culture medium at 500nM per sample for 30 minutes at 37 degrees. TMRM (Invitrogen) staining was performed in culture medium at 5nM per sample for 15 minutes at 37 degrees. BrdU analysis was performed utilizing the Phase-Flow kit per manufacturer’s instructions (BioLegend), with cells cultured in BrDU for 2 hours prior to staining. Flow data was captured on a BD Fortessa analyzer (BD Biosciences) and evaluated using FlowJo software (version 10.1, Tree Star). Cells were gated by forward and side scatter to identify lymphocyte population, then gated on GFP+ or CD34+ (RQR8 marker) cells for downstream analysis.

### Seahorse Mito Stress Assay

The Seahorse XF Cell Mito Stress Test Kit (Agilent, Santa Clara, CA; Catalog #103015-100) was run on a Seahorse XFe96 Bioanalyzer (Agilent) to determine basal and maximal oxygen consumption rates (OCR), spare respiratory capacity (SRC), and extracellular acidification rate (ECAR) for transduced T cells. T cells were plated in assay media (XF Base media (Agilent) with glucose (25mM), sodium pyruvate (2□mM) and L-glutamine (4□mM) (Gibco), pH 7.4 at 37□°C) on a Seahorse cell culture plate coated with Cell-Tak (Corning) at 1×10^5^ cells/well. After adherence and equilibration, basal ECAR and OCR readings were taken for 30 min. Cells were then stimulated with oligomycin (2 μM), carbonyl cyanide 4- (trifluoromethoxy) phenylhydrazone (FCCP, 1 μM), and rotenone/antimycin A (0.5 μM) to obtain maximal respiratory and control values. Assay parameters were as follows: 3□min mix, no wait, 3□min measurement, repeated for 3 cycles at baseline and after each injection. SRC was calculated as the difference between basal and maximal OCR values obtained after FCCP uncoupling. The XF Mito Stress Test report generator and Agilent Seahorse analytics were used to calculate parameters from Wave software (Agilent, Version 2.6.1.53).

### Cytokine multiplex and ELISA analysis

Supernatants were assessed for cytokine production using the LEGENDplex™ Human Inflammation Panel 1 (13-plex) in a V-bottom plate per the manufacturer’s instructions (BioLegend, San Diego, CA, USA). Flow cytometry data were acquired on a BD Fortessa analyzer (BD Biosciences, Franklin Lakes, NJ, USA) and assessed using FlowJo software (version 10.7, Tree Star). Where indicated, enzyme linked immunosorbent assay (ELISA) to detect human IL-8 was done using culture supernatants at 1:100 dilution in a Quantikine ELISA kit (R&D Systems) per manufacturer’s instructions. One sample run by LEGENDplex did not have sufficient volume to be further assessed by ELISA.

### Mixed Leukocyte Reaction

On day 9 of culture, cells were flow sorted for GFP positivity, labeled with CellTrace Violet (Invitrogen), and placed at 3×10^5^/well in culture with 3×10^5^allogeneic non-T cell APCs irradiated at 20Gy. MLRs were performed in 96-well round-bottom plates in RPMI with varying levels of glucose. Physiologic glucose media was obtained by mixing no glucose RPMI (Gibco Cat #11879-020) 1:1 with standard RPMI media (Gibco Cat #11875-085) for a final glucose concentration of 1g/L (5.5mM). Low glucose RPMI was obtained by mixing no glucose RPMI 1:3 with standard RPMI media for a final glucose concentration of 0.5g/L (2.25mM). After 3-4 days, media was collected for cytokine analysis and cells were assessed by flow cytometry, with responding cells identified by decreased levels of CellTrace Violet.

### Statistics

Graphing and statistical analysis was performed using GraphPad Prism for Windows (version 9.0.1, San Diego, CA; www.graphpad.com). Paired two-tailed Student’s t test or one-way analysis of variance (ANOVA) were used to determine statistical significance. All samples with variability between data points were run through a two-sided Grubbs’ test for outliers using GraphPad Prism software. After Grubbs’ analysis, one MCP-1 sample was eliminated as an outlier (making 3 independent donors instead of 4). Unless noted otherwise, data are displayed as mean ± standard error of the mean. In all cases *p<0.05, **p<0.01, ***p<0.001, and ****p<0.0001.

### Study Approval

All studies on human cells were designated Exempt status by the University of Pittsburgh Institutional Review Board.

## Footnotes

This work was supported by grants to CAB from the Department of Defense (CA180681), National Institute of Health – NHBLI (R01 HL144556), the Hyundai Motor Company (Hope on Wheels Scholar grant), and the American Society of Hematology (Scholar award). ELB has received support from the University of Pittsburgh Cancer Immunology Training Program T32 (5T32CA082084), NICHD K12 Grant (HD052892) and the St. Baldrick’s Foundation Fellowship grant. The University of Pittsburgh holds a Physician-Scientist Institutional Award from the Burroughs Wellcome Fund (ELB).

## Notes

Conflict of interest: The authors have declared that no conflict of interest exists.

### Competing Interest Statement

The authors have declared no competing interest.

