## Supplemental Figure 1 for "Overexpression of AMPKγ2 increases AMPK signaling to augment human T cell metabolism and function"

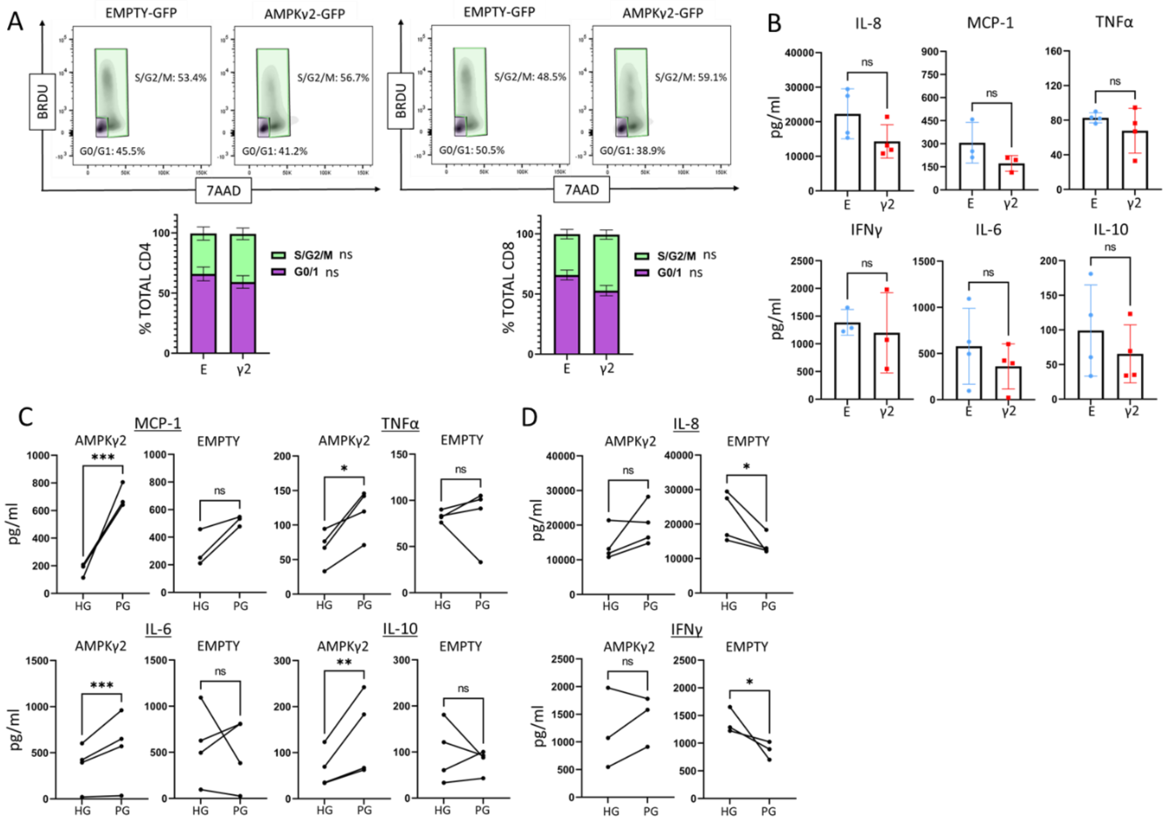


**Supplemental Figure 1. AMPKγ2 transduced cells function similarly to control cells in high glucose media but increase cytokine production when challenged in physiologic glucose. (A)** AMPKγ2- and Empty-transduced T cells were cultured for 9 days then re-stimulated in RPMI (11 mM glucose) for 72 hours, incubated with BrdU for 2 hours, harvested, and counter-stained with 7AAD. **(B-D)** AMPK- and Empty-transduced T cells were cultured for 9 days then plated against allogeneic non-T cell APCs in RPMI containing either 11 mM (high glucose (HG)) or 5.5 mM (physiologic glucose (PG)) for 72 hours. Media was harvested and assessed for cytokine expression by LEGENDplex analysis. Cytokine levels were first compared between AMPK- and Empty-transduced T cells in HG media **(B)** and then in the same cells in HG versus PG **(C-D)**. Compared to HG media, AMPKg2-transduced cells stimulated in PG produced either higher concentrations of inflammatory cytokines while Empty-transduced control cells did not **(C)** or maintained cytokine concentrations that decreased in control cultures **(D).** Flow plots represent data from 2-4 independent donors, while graphs are composites of all data. Bar graphs in **(A)** represent 3 independent donors. $*p<0.05$, $**p<0.01$, $***p<0.001$ by paired Student's T test

### Supplemental Tables

**Supplemental Table 1 – Antibodies for Immunoblot analysis**

| Antigen | Company | Clone | Catalog # |
|---|---|---|---|
| **Phospho-ACC** | *Cell Signaling* | D7D11 | 11818S |
| **ACC** | *Cell Signaling* | C83B10 | 3676S |
| **Phospho-ULK-1** | *Cell Signaling* | D1H4 | 5869S |
| **ULK-1** | *Cell Signaling* | D8H5 | 8054S |
| **Opa1** | *Cell Signaling* | D6U6N | 80471S |
| **Mfn1** | *Cell Signaling* | D6E2S | 14739S |
| **Phospho-AMPK** | *Cell Signaling* | T172 | 2535S |
| **AMPK** | *Cell Signaling* | F6 | 2793S |
| **Beta Actin** | *Cell Signaling* | 13E5 | 4970S |
| **PGC1a** | *Cell Signaling* | 3G6 | 2178S |

Abbreviations: ACC Acetyl CoA Carboxylase, AMPK AMP-activated protein kinase, ULK1 Unc51-like kinase 1

**Supplemental Table 2 – Antibodies and reagents for flow cytometry**

| Antibody | Conjugate | Company | clone |
|---|---|---|---|
| PD1 | Per-CP | BIOLEGEND | EH12.2H7 |
| CD8 | EF780 | INVITROGEN | SK1 |
| LAG3 | PE | BIOLEGEND | 11C3C65 |
| CD4 | PeCY7, APC, PE | INVITROGEN | RPA-T4 |
| CD25 | BV711 | BIOLEGEND | BC96 |
| TIM3 | BV605 | BIOLEGEND | F38-2E2 |
| CD45RA | APC | INVITROGEN | HI100 |
| CD62L | BV605 | BIOLEGEND | DREG-56 |
| IFNG | EF780 | INVITROGEN | 4SB3 |
| IL10 | PeCY7 | INVITROGEN | JES3-9D7 |
| IL2 | BV711 | BD BIOSCIENCES | 5344.111 |
| TNFα | BV605 | BIOLEGEND | MAb11 |
| PS6 | Per-CP | INVITROGEN | cupk43k |
| CD4 | Pac Blue | BIOLEGEND | SK3 |
| CD8 | BV711 | BD BIOSCIENCES | RPA-T4 |
| CD8 | Per-CP | BD BIOSCIENCES | SK1 |
| CD4 | APC | INVITROGEN | RPA-T4 |
| P4EBP1 | PE | INVITROGEN | V3NTY24 |

| Reagent | Company | Cat. No: |
|---|---|---|
| **LIVE/DEAD™ Fixable Dead Cell Stain** | Thermo Scientific | L34957 |
| **MitoTracker Deep Red** | Thermo Scientific | M22426 |
